# Woronin bodies move dynamically and bidirectionally by hitchhiking on early endosomes in *Aspergillus nidulans*

**DOI:** 10.1101/2023.01.20.524968

**Authors:** Livia D. Songster, Devahuti Bhuyan, Jenna R. Christensen, Samara L. Reck-Peterson

## Abstract

The proper functioning of organelles depends on their intracellular localization, mediated by motor protein-dependent transport on cytoskeletal tracks. Rather than directly associating with a motor protein, peroxisomes move by hitchhiking on motile early endosomes in the filamentous fungus *Aspergillus nidulans*. However, the cellular function of peroxisome hitchhiking is unclear. Peroxisome hitchhiking requires the protein PxdA, which is conserved within the fungal subphylum Pezizomycotina, but absent from other fungal clades. Woronin bodies are specialized peroxisomes that are also unique to the Pezizomycotina. In these fungi, multinucleate hyphal segments are separated by incomplete cell walls called septa that possess a central pore enabling cytoplasmic exchange. Upon damage to a hyphal segment, Woronin bodies plug septal pores to prevent wide-spread leakage. Here, we tested if peroxisome hitchhiking is important for Woronin body motility, distribution, and function in *A. nidulans*. We show that Woronin body proteins are present within all motile peroxisomes and hitchhike on PxdA-labeled early endosomes during bidirectional, long-distance movements. Loss of peroxisome hitchhiking by knocking out *pxdA* significantly affected Woronin body distribution and motility in the cytoplasm, but Woronin body hitchhiking is ultimately dispensable for septal localization and plugging.

## Introduction

Within a eukaryotic cell, organelles are precisely positioned to regulate their cellular function. The intracellular trafficking of these cargos is primarily achieved by molecular motors that bind different cargos and directly transport them along cytoskeletal filaments like microtubules or actin^1,2^. Microtubules are polar structures that serve as the primary track for transporting cargo long distances in eukaryotes. They possess a fast-growing plus end typically oriented towards the cell periphery and a minus end originating at a microtubule organizing center. Kinesin motors transport cargo predominantly towards the plus end of microtubules (anterograde)^3^. Conversely, cytoplasmic dynein motors (“dynein” here) transport cargo toward microtubule minus ends (retrograde)^1^.

The canonical view of intracellular transport is that a cargo attaches directly to a motor protein for movement, typically via a protein adaptor^1,4^. However, a novel form of intracellular transport termed “hitchhiking” was recently discovered, whereby a primary cargo directly interacts with a motor and a secondary hitchhiking cargo is transported indirectly via association with the primary cargo (review^5^). Hitchhiking-like phenomena have been observed in diverse cell types and organisms such as filamentous fungi^6,7^, animal neurons^8,9^, and plant cells^10,11^.

In *Aspergillus nidulans*, peroxisomes move long distances along microtubules by hitchhiking on motile early endosomes transported by kinesin-3 or dynein^7^. We previously identified a putative molecular tether (<u>p</u>eroxisome <u>d</u>istribution <u>m</u>utant A; PxdA) and a phosphatase (<u>D</u>enA-interacting <u>p</u>hosphatase A; DipA) that are essential for peroxisome hitchhiking on early endosomes^7,12^. Both PxdA and DipA are found on early endosomes and loss of either compromises long distance motility of peroxisomes, resulting in an accumulation of peroxisomes in the hyphal tip without affecting early endosome motility or distribution^7,12^. Peroxisomes are the only organelle demonstrated to hitchhike on early endosomes in *A. nidulans*^12^, but the cellular basis for why peroxisomes hitchhike is unclear.

Using comparative genomics, we found that PxdA is conserved within the Pezizomycotina subphylum of ascomycete filamentous fungi, suggesting that a function for peroxisome hitchhiking might also be conserved within this clade. We therefore investigated if hitchhiking was required for septal plugging, a peroxisome function unique to the Pezizomycotina.

Septal plugging is a critical process within Pezizomycotina fungi as they possess continuous hyphal compartments separated only by a septal wall containing a pore^13^. This interconnected hyphal network or “mycelium” has the advantage of quickly exchanging components throughout the organism^14,15^. However, if one portion of the organism is injured or lysed, significant cytoplasmic leakage can occur. In many species within the Pezizomycotina, peroxisome derivatives known as Woronin bodies plug septal pores upon mycelial injury, allowing the rest of the mycelium to remain intact^16^. Woronin bodies can also function to dynamically open and close pores to control hyphal heterogeneity^14^. Woronin body biogenesis is thought to initiate at the growing hyphal tip, based on a spatial bias in mRNA expression of the essential Woronin body component Hex-1 in *Neurospora crassa* ^17^. After biogenesis at the hyphal tip, Woronin bodies are thought to be transported to nascent septal sites, where they bud from peroxisomes and remain tethered at the septa^18,19^. However, the mechanism by which Woronin bodies are transported to septa is unknown.

In this study, we investigated whether peroxisome hitchhiking is important for Woronin body motility, positioning, and function in *A. nidulans*. We find that Woronin body proteins associated with peroxisomes move dynamically and bidirectionally, and that this motility is dependent upon PxdA-mediated hitchhiking on early endosomes. We show that Woronin bodies separate from peroxisomes once localized to septa, but this fission process does not require PxdA. Finally, we show that the distribution of Woronin bodies in hyphal tips requires hitchhiking, but this is ultimately dispensable for the positioning of Woronin bodies at septa and their function in septal closure after hyphal injury.

## Results

### Peroxisome hitchhiking proteins and Woronin body specialization are conserved within the Pezizomycotina fungi

Peroxisome hitchhiking has been directly observed in two species of fungi, *A. nidulans* and *Ustilago maydis*^6,7^. However, a cellular function for peroxisome hitchhiking has not been established. Two early endosome-associated proteins, a putative molecular tether (PxdA) and a phosphatase (DipA), have been shown to be essential for peroxisome hitchhiking in *A. nidulans* (Fig. 1A-B)^7,12^.

**Figure 1.**
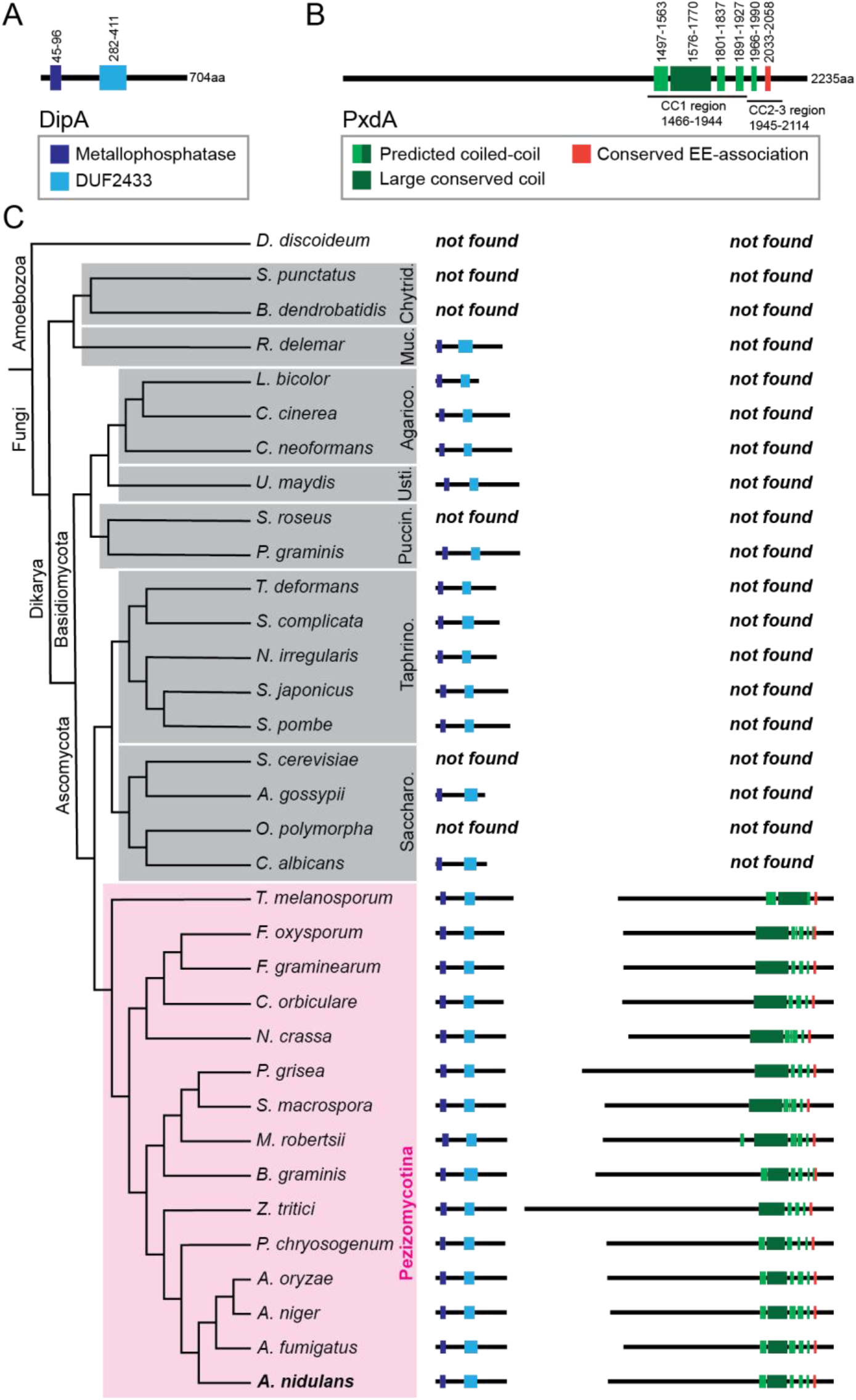
Phylogenetic analysis of the essential peroxisome hitchhiking proteins DipA and PxdA. **(A-B)** Protein domain cartoons for *A. nidulans* DipA (A) and PxdA (B). The domains previously defined^7^ as CC1 and CC2-3 are indicated. Green bars represent coiled coils predicted in this study. **(C)** Phylogenetic tree showing conservation of DipA and PxdA across 33 species representing major fungal clades, indicated by gray and pink boxes (clade abbreviations: Saccharo. = Saccharomycotina; Taphrino = Taphrinomycotina; Puccin. = Pucciniomycotina; Usti. = Ustilagomycotina; Agarico. = Agaricomycotina; Muc. = Mucoromycotina; Chytrid. = Chytridiomycota). *Dictyostelium discoideum* was included as an outgroup. Protein domain cartoons are scaled to one another to display amino acid length and domain organization in different species.

To search for a potential cellular role for peroxisome hitchhiking, we used comparative genomics to identify which fungal clades likely exhibit PxdA-dependent peroxisome hitchhiking. We used BLASTp to search for DipA and PxdA protein homologs in 33 fungal species representing distinct clades across the fungal tree, with the underlying assumption that fungal species encoding both PxdA and DipA in their genomes would exhibit peroxisome hitchhiking. Putative PxdA homologs were only identified in the Pezizomycotina clade and were present in all 15 Pezizomycotina species we analyzed (Fig. 1C; Table S1).

Peroxisome hitchhiking has also been observed outside of the Pezizomycotina, in the basidiomycete *U. maydis*^6^. We did not identify a homolog of PxdA in *U. maydis*, suggesting that peroxisome hitchhiking may function by a distinct mechanism in this species. However, it is possible that *U. maydis* possesses a PxdA homolog that is sequence divergent and therefore not identified by our methods.

Putative DipA homologs were identified in all queried Pezizomycotina species as well as in other fungal clades (Table S1). DipA functions in cellular processes beyond peroxisome hitchhiking, including septal positioning^20^, likely explaining why DipA is found more broadly in the fungal kingdom.

As putative homologs for both PxdA and DipA were found only in the Pezizomycotina, we hypothesized that a cellular function for peroxisome hitchhiking may also be specific to these fungi. The plugging of septal pores by Woronin bodies, which are derived from peroxisomes, also occurs exclusively within the Pezizomycotina ascomycetes^21^. Additionally, previous work has suggested that Woronin bodies might require active transport to reach nascent septal assembly sites prior to budding from peroxisomes^19^. We therefore sought to test if peroxisome hitchhiking affected Woronin body localization and function in septal pore plugging in *A. nidulans*.

### Characterization of Woronin body biogenesis and distribution in *Aspergillus nidulans*

To determine if Woronin body function requires hitch-hiking, we first needed to reliably distinguish peroxisomes and Woronin bodies from one another in live cells. We achieved this by quantifying the colocalization and co-occurrence of fluorescently tagged proteins that were either an essential Woronin body component or a peroxisome marker. We chose SspA and HexA as representative Woronin body proteins for *A. nidulans* based on their involvement in Woronin body formation and function, which has been most extensively examined in *N. crassa*, another member of the Pezizomycotina (Fig. 1C; Table S1).

In *N. crassa*, the proteins WSC (<u>W</u>oronin <u>s</u>orting <u>c</u>omplex; SspA in *A. nidulans*) and Hex-1 (HexA in *A. nidulans*) are essential for Woronin body formation and function, respectively. Previous work has shown that peroxisomes begin developing Woronin bodies with the import of Hex-1 via its C-terminal peroxisome targeting signal (PTS1)^16^. After import, Hex-1 self-assembles into a dense crystalline core, the main structural component of the Woronin body^16,22^. The Hex-1 complex then polarizes to one side of the peroxisome due to the presence of the WSC protein in the peroxisome membrane^23^. This polarization is followed by organelle fission and separation of the Woronin body from the peroxisome^17,24^.

The formation of Woronin bodies in *A. nidulans* is largely similar to that of *N. crassa*, as they also mature from peroxisomes following the import of HexA and SspA (Fig. 2A)^25,26^. To characterize the colocalization of Woronin bodies with peroxisomes in *A. nidulans*, we quantified the degree of overlap between endogenously tagged HexA, SspA, and the peroxisome marker PTS1 via fluorescence microscopy in germlings. We found almost all SspA puncta had co-occurring HexA signal. On the other hand, a large population of HexA puncta had no corresponding SspA signal (Fig. S1A-B). PTS1 co-occurred more frequently with HexA (Fig. S1C-D) than it did with SspA (Fig. S1E-F), suggesting that HexA is imported into peroxisomes prior to SspA. Consistent with this finding, PTS1 and HexA were often, but not always, closely overlapping (Fig. S1C), whereas PTS1 and SspA were often adjacent and only weakly overlapping with one another (Fig. S1E). Together, these data suggest that while the majority of peroxisome and Woronin body proteins exist within the same membrane-bound compartment, distinct stages of Woronin body maturation can be visualized using fluorescent microscopy.

**Figure 2.**
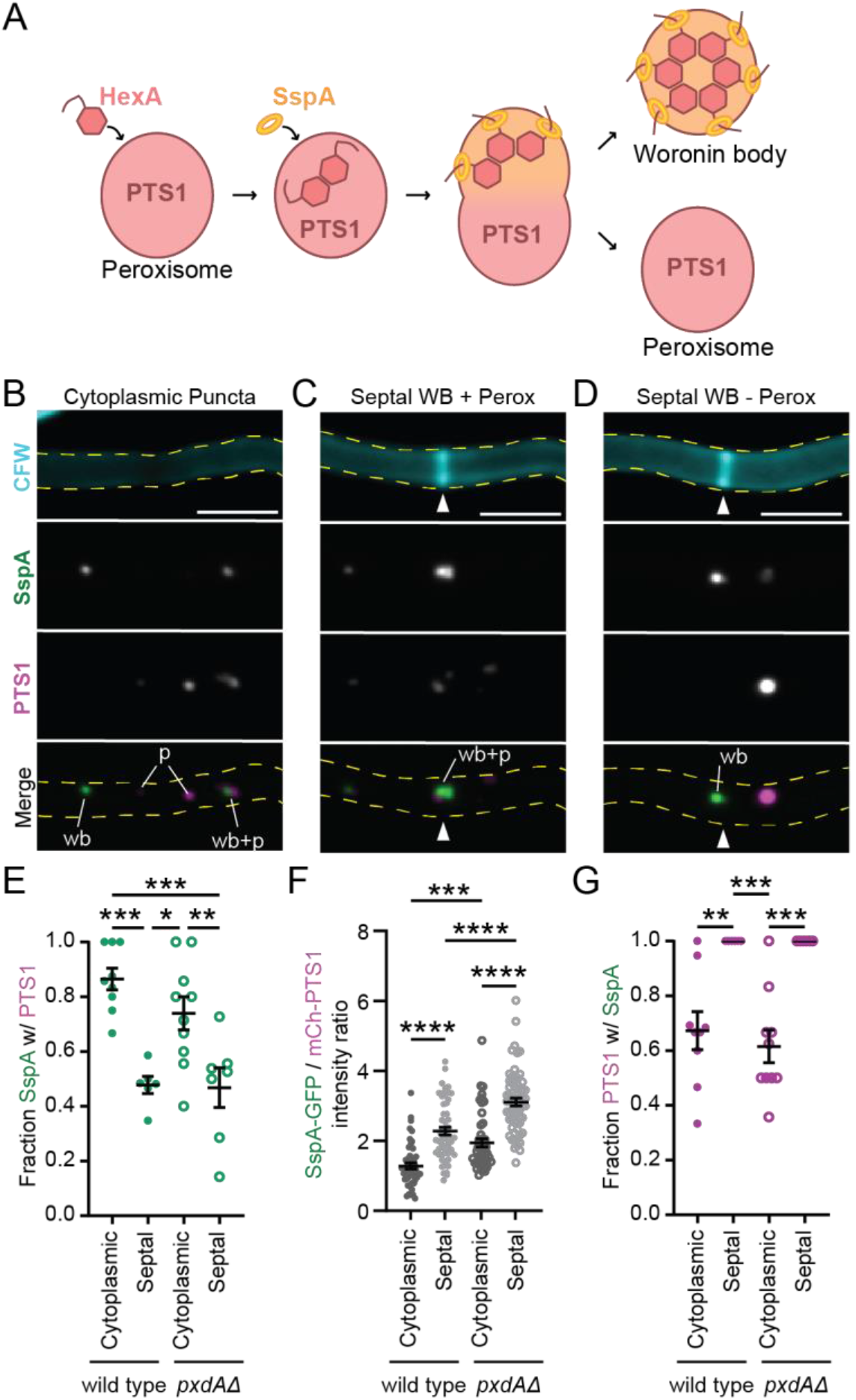
Characterization of Woronin body maturation from peroxisomes in the cytoplasm versus at septa. **(A)** Diagram of Woronin body maturation from peroxisomes after import of HexA and SspA from the cytoplasm. **(B-D)** Example of SspA-mGFP5 and mCherry-PTS1 in the cytoplasm (B-C) and at tip apical septa (D) of adult wild type hyphae (sum-Z projections). White triangles indicate septa. CFW = calcofluor white, cell membrane dye; wb = Woronin body; p = peroxisome. Scale bars = 5 μm. **(E)** Average SspA-mGFP5 co-occurrence with mCherry-PTS1 while in the cytoplasm or associated with septa. Each dot represents one experiment with 5 to 20 regions of interest scored (n fields of view, mean ± SD: wild type cytoplasmic = 9, 0.87 ± 0.12; wild type septal = 6, 0.48 ± 0.08; *pxdAΔ* cytoplasmic = 10, 0.74 ± 0.19; *pxdAΔ* septal = 7, 0.47 ± 0.019). **(F)** Average intensity ratio of SspA-mGFP5 and mCherry-PTS1 in puncta at septa and in the cytoplasm (n puncta, mean ± SD: wild type cytoplasmic = 49, 1.28 ± 0.63; wild type septal = 56, 2.28 ± 0.84; *pxdAΔ* cytoplasmic = 48, 1.94 ± 0.82; *pxdAΔ* septal = 62, 3.11 ± 0.89). **(G)** Average mCherry-PTS1 co-occurrence with SspA-mGFP5 in the cytoplasm or associated with septa. Each dot represents one experiment with 5 to 20 regions of interest scored (n fields of view, mean ± SD: wild type cytoplasmic = 9, 0.67 ± 0.21; wild type septal = 6, 1.00 ± 0; *pxdAΔ* cytoplasmic = 10, 0.62 ± 0.19; *pxdAΔ* septal = 7, 1.00 ± 0). **For E-G:** Ordinary one-way ANOVA with post-hoc Tukey-Kramer test for multiple comparisons. Error bars represent SEM. P > 0.05 (labeled ns for “not significant,” or not shown), P ≤ 0.05 (*), P ≤ 0.01 (**), P ≤ 0.001 (***), P ≤ 0.0001 (****).

These colocalization experiments in live cells established that SspA could serve as a sufficient marker for later stages of Woronin body maturation. We therefore used this as a tool to visually distinguish how Woronin body (SspA-mGFP5) and peroxisome (mCherry-PTS1) colocalization varies at different intracellular positions in wild type hyphae (Fig. 2A). We began by scoring the fraction of colocalized Woronin body and peroxisome puncta in the cytoplasm (Fig. 2B) versus at septa (Fig. 2C-D). Cytoplasmic Woronin bodies were frequently associated with a peroxisome in wild type hyphae (Fig. 2E) and rarely found without an attached peroxisome (Fig. 2B). In comparison, at septa, about half of Woronin bodies had no associated peroxisome signal (Fig. 2D-E). For those septal Woronin bodies with corresponding peroxisome signal, the Woronin body signal was always closer to the septa (Fig. 2C; white arrow). This preferred orientation is likely due to Woronin body tethering at septal sites by HexA^27^.

To better measure the relative amounts of peroxisome versus Woronin body proteins in the cytoplasm and at septa, we quantified the fluorescent intensity of SspA and HexA relative to PTS1. The ratio of SspA to HexA was consistent at both septa and in the cytoplasm (Fig. S1G), suggesting Woronin body composition does not vary much between both locations. In contrast, septal puncta exhibited a significantly higher intensity ratio of SspA to PTS1 than cytoplasmic puncta (Fig. 2F). We observed a similar pattern for the ratio of HexA to PTS1 (Fig. S1H). Together, this data supports the idea that most Woronin bodies have separated from peroxisomes by the time they associate with septa.

Finally, we wanted to estimate how many cytoplasmic peroxisomes were actively developing Woronin bodies. We observed that 67% of all cytoplasmic peroxisomes had an associated Woronin body (Fig. 2G). Overall, these data show that not all peroxisomes in the cytoplasm are actively developing Woronin bodies in *A. nidulans*. However, for peroxisomes that have begun developing Woronin bodies, the two organelles remain attached in the cytoplasm, and only separate shortly before or immediately following recruitment to septa.

Previous work has established that in most Pezizomycotina including *A. nidulans*, Woronin bodies are tethered at septa^18^. However, a small number of species including *N. crassa* use a divergent mechanism of Woronin body tethering, whereby Woronin bodies are delocalized from septa and instead are tethered to the actin cortex throughout a hyphal compartment^28^. Our observations in *A. nidulans* on the localization of Woronin body maturation and separation from peroxisomes after septal association is similar to what was previously reported for *N. crassa*^28^, despite these species exhibiting different Woronin body localization and tethering mechanisms.

### Woronin bodies move bidirectionally using the peroxisome hitchhiking protein PxdA

Woronin body biogenesis is thought to occur primarily in the cytoplasm near the hyphal tip^17,18^. Woronin bodies are then localized to their tethering site at or near septal pores, where they function to seal septal pores following injury^19^. However, the mechanism by which they are transported to septa is unknown. We therefore wanted to characterize Woronin body movement in hyphae and determine if peroxisome hitchhiking was responsible for Woronin body movement and transport to septa.

We showed that Woronin bodies frequently colocalize with peroxisomes in the cytoplasm but are present alone at septa (Fig. 2E). This suggests that Woronin bodies could either (1) move in conjunction with hitchhiking peroxisomes and bud off once present at septa or (2) bud off peroxisomes in the cytoplasm and be transported to septa by a distinct mechanism. To test this, we first compared Woronin body and peroxisome movement in adult hyphae. In *A. nidulans*, only a small population of peroxisomes exhibit long-distance movements at a given time^7,29^. We found that Woronin bodies also exhibit infrequent but dynamic bidirectional movements, like peroxisomes (Movie S1). As expected^7^, 10% of total movements for both peroxisomes and Woronin body movements were processive over long-distances (>3 μm), indicative of microtubule-driven movement. During these long-distance movements, Woronin bodies moved at similar speeds (Fig. 3A) and over similar distances to peroxisomes (Fig. 3B). Both peroxisomes and Woronin bodies also exhibited bidirectional movements with no directional bias (Fig. 3C).

**Figure 3.**
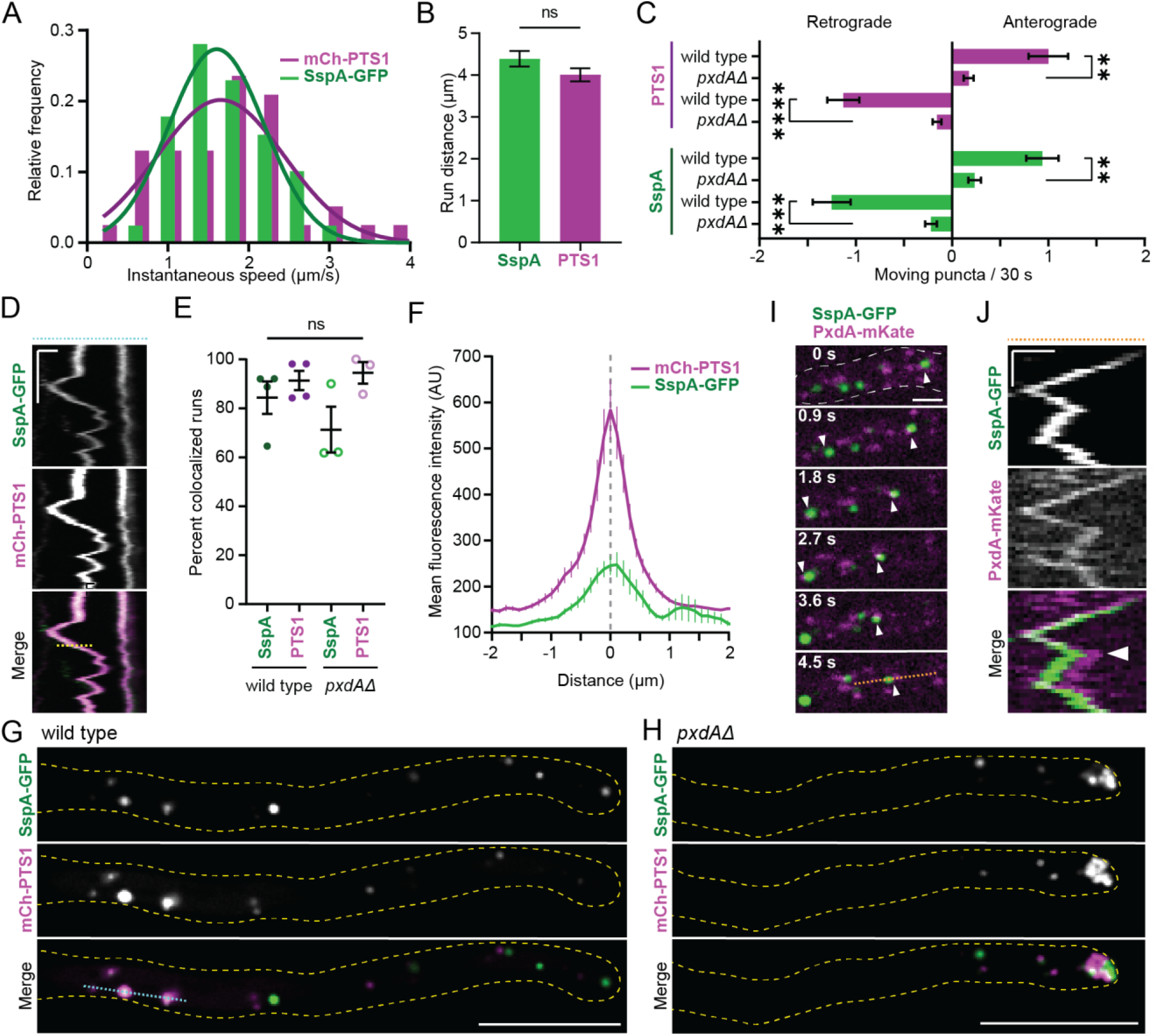
Woronin bodies at the hyphal tip exhibit dynamic, bidirectional movement with peroxisomes, and this movement requires PxdA. **(A)** Histogram of speeds of long-distance runs (>3 μm travelled in one direction) for SspA-mGFP5 and mCherry-PTS1 puncta (n directed runs/total, mean ± SD: SspA-mGFP5 = 39/390, 1.71 ± 0.59 μm/s; mCherry-PTS1 = 34/330, 1.72 ± 0.79 μm/s). Solid lines indicate nonlinear Gaussian fit lines (SspA-mGFP5: R^2^ = 0.95; mCherry-PTS1: R^2^ = 0.76). **(B)** Mean distance of long-distance runs for SspA-mGFP5 and mCherry-PTS1 (n, mean ± SD: SspA-mGFP5 = 39 runs, 4.39 ± 1.2 μm; mCherry-PTS1 = 34 runs, 4.01 ± 0.96 μm). **(C)** Number of anterograde (towards the hyphal tip) and retrograde (towards the apical septum) movements of mCherry-PTS1 (top) and SspA-mGFP5 (bottom) in wild type and *pxdAΔ* adult hyphae (n: wild type = 32 cells and *pxdAΔ* = 64 cells. PTS1 mean ± SD puncta/30 s: anterograde wild type = 1.0 ± 1.2; anterograde *pxdAΔ* = 0.17 ± 0.42; retrograde wild type = -1.1 ± 0.94; retrograde *pxdAΔ* = -0.16 ± 0.37. SspA mean ± SD: anterograde wild type = 0.94 ± 0.95; anterograde *pxdAΔ* = 0.23 ± 0.53; retrograde wild type = -1.25 ± 1.1; retrograde *pxdAΔ* = -0.19 ± 0.50). **(D)** Kymograph depicting mCherry-PTS1 and SspA-mGFP5 colocalization over time. This kymograph was generated by the blue dashed line shown in (G). The yellow dashed line on the kymograph is an example of a line used for (F). X-scale bar = 1 μm, Y-scale bar = 10 s. **(E)** Percent of long-distance runs (>3 μm) that are colocalized; each dot represents an individual experiment, where all runs from at least 10 hyphal tips were scored for colocalization. The number of moving SspA puncta ranged from 11-18 for *pxdAΔ* and from 14-30 for wild type experiments. The number of moving PTS1 puncta ranged from 6-13 for *pxdAΔ* and from 7-26 for wild type experiments (n experiments, mean ± SD: wild type SspA = 4, 0.84 ± 0.13; wild type PTS1 = 4, 0.91 ± 0.08; *pxdAΔ* SspA = 3, 0.71 ± 0.16; *pxdAΔ* PTS1 = 3, 0.95 ± 0.08). **(F)** Mean intensity of SspA-mGFP5 and mCherry-PTS1 during comigrating runs in wild type (n = 53 runs collected from 23 cells). The gray dashed line indicates the middle of the mCherry-PTS1 peak. **(G-H)** SspA-mGFP5 and mCherry-PTS1 in a single plane of wild type (G) and *pxdAΔ* (H) hyphal tips. The blue dashed line was used to generate kymograph in (D). Scale bars = 10 μm. **(I)** SspA-mGFP5 and PxdA-mKate comigration towards a growing hyphal tip (not shown; to the left). White arrows indicate two comigration events. The dashed orange line in the last panel was used to generate the kymograph in (I). Scale bar = 2 μm. **(J)** Kymograph depicting SspA-mGFP5 and PxdA-mKate colocalization over time. X-scale bar = 1 μm, Y-scale bar = 3 s. White arrow indicates the time point at around 6 s where PxdA-mKate switches from leading SspA-mGFP5 to following. **For B-C and E:** Error bars represent SEM. Unpaired t-test; P > 0.05 (labeled ns for “not significant,” or not shown), P ≤ 0.05 (*), P ≤ 0.01 (**), P ≤ 0.001 (***), P ≤ 0.0001 (****).

Next, we sought to test how frequently Woronin bodies and peroxisomes colocalize during long-distance runs (>3 μm travelled in one direction). The large majority of moving Woronin bodies were colocalized with a peroxisome, and vice versa (Fig. 3D-E). This finding was surprising as we previously observed that only two thirds of cytoplasmic peroxisomes had associated Woronin body signal (Fig. 2G). Furthermore, we found no obvious bias in whether the Woronin body was ‘leading’ or ‘trailing’ the peroxisome during motile runs, and instead observed the signals frequently overlapped during movement (Fig. 3F). This contrasts with our observations on stationary cytoplasmic peroxisomes and Woronin bodies, which frequently exhibited Woronin body signal that was polarized and offset to one side of the peroxisome signal (Fig. 2B). Together, this data suggests that Woronin body proteins move via association with a hitchhiking peroxisome, and not as a separate / distinct organelle.

We next tested if Woronin body movement required the essential peroxisome hitchhiking protein PxdA. Compared to wild type hyphae, in which peroxisomes and Woronin bodies are consistently distributed across hyphae (Fig. 3G), *pxdAΔ* hyphae had an accumulation of both peroxisomes and Woronin bodies at the hyphal tip (Fig. 3H) and a drastic reduction in the number of Woronin bodies and peroxisomes moving in either direction (Fig. 3C; Movie S2). Furthermore, motile Woronin bodies colocalized with the putative hitchhiking tether PxdA (Fig. 3I-J). These data demonstrate that Woronin bodies require PxdA for long-distance, bidirectional movements in hyphal tips.

While loss of PxdA abolished most long-distance movements, a small number of peroxisomes and Woronin bodies still moved processively in *pxdAΔ* hyphal tips (Fig. S1K-L). However, these Woronin bodies were still colocalized with peroxisomes during this movement (Fig. 3E). Additionally, the overall Woronin body and peroxisome co-occurrence (SspA/PTS1) in *pxdAΔ* hyphae was similar to what was observed in wild type (Fig. 2E-G, S1F, and S1J). These findings demonstrate that the majority of Woronin body movement occurs via association with hitchhiking peroxisomes and that PxdA is not important for the assembly of Woronin bodies.

### Essential Woronin body proteins are not required for peroxisome hitchhiking

As Woronin bodies and peroxisomes comigrated during long-distance runs, and most motile peroxisomes colocalized with Woronin bodies while moving (Fig. 2E), we wondered if the presence of Woronin body proteins was required for peroxisome motility. We tested this using mutants lacking either HexA or SspA (Fig. S2A). Loss of HexA or SspA did not significantly affect the average number of moving peroxisomes per cell (Fig. S2B). In *hexAΔ* hyphae, peroxisomes showed a slight accumulation at hyphal tips (Fig. S2C), but to a lesser extent than the accumulation observed in *pxdAΔ* hyphae (Fig. S2D). Despite nearly all moving peroxisomes possessing Woronin body proteins, these data show that the ability to form Woronin bodies is not required for peroxisome hitchhiking.

### Woronin bodies localize at septa and plug damaged hyphae independently of hitchhiking

We next wanted to test whether peroxisome-Woronin body hitchhiking is required for Woronin bodies to localize to septa in undamaged hyphae. First, we quantified the cytoplasmic localization of Woronin bodies in strains lacking peroxisome movement (*pxdAΔ* and *hookAΔ*) as well as in a strain with non-functional Woronin bodies (*hexAΔ*). We included *hookAΔ* because HookA is the adaptor protein that links microtubule motors to early endosomes in *A. nidulans* ^30^, and loss of HookA therefore disrupts both early endosome and peroxisome distribution and motility^7^. As predicted, loss of peroxisome movement in both *pxdAΔ* and *hookAΔ* hyphae results in an accumulation of SspA-labeled Woronin bodies at the hyphal tip (Fig. 4A-B). There were occasionally very bright Woronin bodies at the apex of wild type hyphal tips, corresponding to a slight peak in intensity around 0.5 μm from the tip (Fig. 4B, black line) that was abolished in *hexAΔ* hyphae (Fig. 4B, red line). This peak in SspA intensity in wild type tips could correspond to higher levels of SspA import into peroxisomes in this region, or the presence of larger Woronin body/peroxisome organelles that have not yet undergone fission.

**Figure 4.**
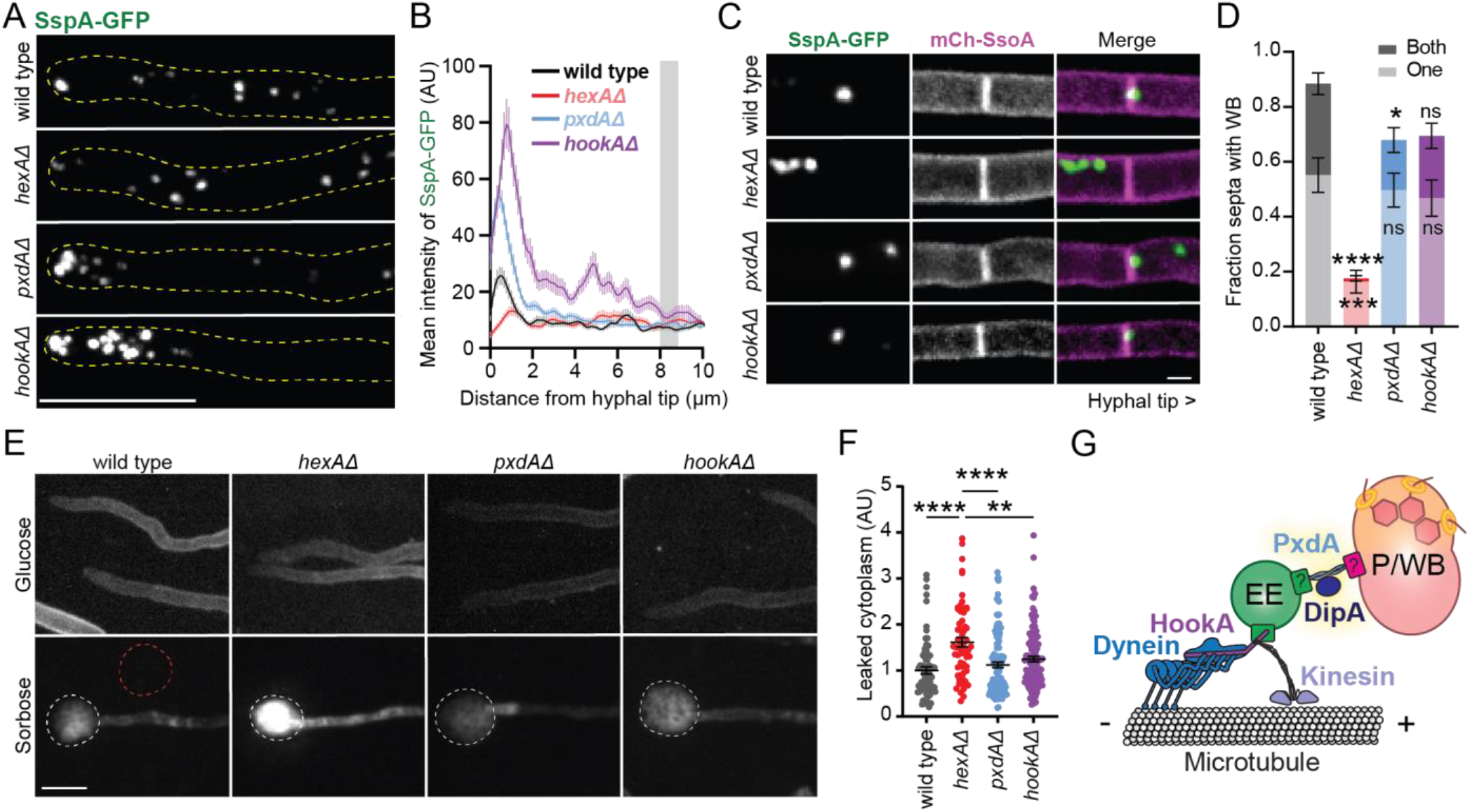
Woronin bodies require hitchhiking to localize at septa, but this localization is not required for septal plugging after hyphal damage. **(A)** SspA-mGFP5 localization at the hyphal tip in adult hyphae (max-Z projections). Scale bar = 10 μm. **(B)** Mean intensity of SspA-mGFP5 from the hyphal tip inwards (n hyphae: wild type = 140; *hexAΔ* = 92; *pxdAΔ* = 225; *hookAΔ* = 132). Gray bar indicates approximate position of a tip apical nucleus. **(C)** Association of Woronin bodies at tip apical septa when grown in glucose MM (max-Z projections). Hyphal tips would be to the right of each image. Scale bar = 2 μm. **(D)** Fraction apical septa with Woronin bodies, quantified as being on one side (lighter bars; bottom) or both sides (darker bars; top) of septa. Each experiment scored 8-64 apical septa (n experiments, mean ± SD: one side wild type = 7, 0.55 ± 0.17; one side *hexAΔ* = 8, 0.16 ± 0.12; one side *pxdAΔ* = 6, 0.50 ± 0.15; one side *hookAΔ* = 4, 0.47 ± 0.13; both sides wild type = 7, 0.32 ± 0.12; both sides *hexAΔ* = 8, 0.012 ± 0.027; both sides *pxdAΔ* = 6, 0.18 ± 0.11; both sides *hookAΔ* = 4, 0.23 ± 0.092). **(E)** Hyphae expressing mCherry-SsoA as a cell membrane marker, when grown in 1% glucose agar versus 2% sorbose/0.1% glucose agar (sum-Z projections). Scale bar = 10 μm. The white circles indicate leaked cytoplasm due to hyphal tip bursting, and the red circle is an example of a background fluorescence measurements. **(F)** Leaked cytoplasm, measured as the integrated density – (background * area) to correct for background intensity, and then normalized to the average wild type (n burst cells, mean ± SD: wild type = 73, 1.00 ± 0.64; *hexAΔ* = 63, 1.61 ± 0.81; *pxdAΔ* = 125, 1.12 ± 0.67; *hookAΔ* = 119, 1.25 ± 0.65). **(G)** Model of peroxisome (P) and Woronin body (WB) hitchhiking on early endosomes (EE). Dynein refers to dynein and dynactin complexes. **For D and F:** Error bars represent SEM. Ordinary one-way ANOVA with post-hoc Tukey-Kramer test; P > 0.05 (labeled ns for “not significant”), P ≤ 0.05 (*), P ≤ 0.01 (**), P ≤ 0.001 (***), P ≤ 0.0001

We next examined Woronin body localization at tip apical septa in undamaged, wild type hyphae as well as *pxdAΔ, hookAΔ, and hexAΔ* mutants. We chose tip apical septa because these septa were most recently formed ^31^ and are still capable of dynamically opening and closing via septal plugging^14^. Transmission electron microscopy studies in *A. nidulans* have previously reported between four and nine Woronin bodies at septa^18^. In assessing Woronin body localization at septa via fluorescence microscopy, three classes of Woronin body localization were identified: (i) septa with Woronin body signal on either side likely representing more than one Woronin body, (ii) septa with Woronin body signal on only one side and (iii) no Woronin body signal (Fig. S3A-B).

Manual scoring of these localization patterns revealed that most tip apical septa had Woronin bodies on one or both sides in wild type hyphae (Fig. 4C-D). Furthermore, approximately one third of tip apical septa had Woronin bodies on both sides in wild type (Fig. 4D). As expected, most *hexAΔ* hyphae lacked Woronin bodies at apical septa. Both *pxdAΔ* and *hookAΔ* hyphae had an intermediate phenotype, with a slight reduction in the number of septa with Woronin bodies on both sides, but similar numbers of septa with Woronin bodies on one side compared to wild type. This finding suggests that PxdA-mediated hitchhiking could contribute to achieving Woronin bodies on both sides of nascent septa. However, fluorescence measurements show there is no significant difference in SspA signal at septa in wild type versus *pxdAΔ* or *hookAΔ* hyphae (Fig. S3C). Overall, our data suggest that Woronin body localization at septa does not depend on Woronin body hitchhiking. Though Woronin body hitchhiking may have a minor contribution to Woronin body recruitment at septa, it is not the primary mechanism by which these organelles localize at septa. Woronin bodies in *pxdAΔ* hyphae could be potentially recruited to tip-apical septa by other mechanisms, such as cytoplasmic movement via ‘poleward drift’^32^ or passive association with septal assembly sites that are marked as the hyphal tip grows^31^.

Finally, we wanted to test if Woronin body hitchhiking was important for Woronin body-based sealing of septal pores following hyphal damage. Growth on sorbose media induced tip lysis in just under half of cells, regardless of genotype (Fig. S3D). Tip lysis and cytoplasmic puddling was clearly visible on sorbose media for all strains (Fig. 4E). The amount of leaked protein was higher in lysed *hexAΔ* than lysed wild type, *pxdAΔ*, or *hookAΔ* cells (Fig. 4F), despite leakage areas appearing similar in size (Fig. S3E). This pattern of cytoplasmic leakage was recapitulated when dried mycelia were exposed to hypotonic shock via distilled water (Fig. S3F). Together, these data demonstrate that Woronin body hitchhiking is not required for proper septal plugging after hyphal damage.

## Concluding thoughts

This study shows that Woronin bodies move dynamically and bidirectionally in the hyphal tip by hitchhiking on PxdA-labeled early endosomes in *A. nidulans*. Though Woronin bodies have been described to move in some fungi (*Z. tritici*^33^; *A. fumigatus*^19^; *A. nidulans*^26^), their motility has not been well characterized and the mechanism by which they move was previously unknown. We showed that Woronin bodies and peroxisomes often coexist as connected organelles in the cytoplasm, where they hitchhike together on early endosomes via PxdA with no directional bias (Fig. 4G). We have identified that virtually all peroxisomes colocalize with a Woronin body while hitchhiking, despite only two thirds of peroxisomes colocalizing with Woronin bodies while immotile in the cytoplasm. Despite this intriguing finding, we show that Woronin body proteins are not required for the identification of peroxisomes as hitchhiking cargo.

It was previously hypothesized that Woronin body biogenesis begins in the hyphal tip, after which they are transported retrograde towards tethering sites at nascent septa^17–19^. We found that Woronin body movement in hyphal tips is bidirectional and requires the hitchhiking protein PxdA. Regardless, we ultimately determined that Woronin bodies do not require PxdA-mediated hitchhiking to reach nor associate with septa, thus disproving the hypothesis that they are actively transported retrograde towards septa.

## Acknowledgements

LDS is funded by an NSF-GRFP and the UCSD Quantitative Integrative Biology T32 from the NIH (1T32GM127235). JRC is funded by a MOSAIC award (K99/R00) from the NIH (K99GM140269). SLRP is supported by HHMI and NIH (1R35GM141825). We thank John Salogiannis and Valentin Wernet for ideas and insight, Kaeling Tan for cloning RPB564, James Holcomb for cloning RPB2190, and Xin Xiang for the mGFP5 plasmid. This paper was typeset with the bioRxiv word template by @Chrelli: www.github.com/chrelli/bioRxiv-word-template

## Author contributions

LS, JRC, and SLR-P devised the experiments. LS and DB performed the experiments. JRC and SLR-P supervised the research. LS, JRC, and SLR-P wrote and edited the manuscript.

## Materials and Methods

### Strain construction

*A. nidulans* strains used in this study are listed in Table S2. New strains were generated by transforming PCR-amplified targeting DNA into *A. nidulans* protoplasts^34^ or via genetic crosses^35^. The genotypes of novel strains were confirmed by PCR amplification from isolated genomic DNA^36^ and Sanger sequencing.

### Plasmid cloning

Plasmids and primers generated for this study are listed in Table S3 and Table S4, respectively. All DNA constructs were designed for homologous recombination at the endogenous locus with the endogenous promoter unless otherwise stated. Novel plasmids were cloned using Gibson isothermal assembly^37,38^. Briefly, fragments were amplified using PCR (NEB Q5 highfidelity DNA polymerase, M0491) from other plasmids or *A. nidulans* genomic DNA. Then, fragments were assembled into the Blue Heron Biotechnology pUC vector at the 5′EcoRI and 3′HindIII restriction sites. The cloning of *mTagGFP2-rabA::AfpyrG, mCherry-FLAG-PTS1::AfpyroA*, and *2xBFPPTS1::AfriboB* were described previously^39^. The cloning of *pxdAmKate2::AfpyroA* and *pxdAΔ::AfriboB* were also previously published^7,12^. All fluorescently tagged proteins had a GA5 linker. The sequence for mGFP5 was optimized for expression in *A. thalania* plants^40^.

### Fungal growth conditions

*A. nidulans* strains were grown in either liquid or solid agar containing yeast-glucose (YG) complete media or minimal media (MM)^34^. YG and MM agar plates contained 10 g/L agar-agar/gum agar (USB 10654). YG complete media contained Bacto Yeast Extract (5 g/L), 2% final D+ glucose (20 g/L), trace elements solution (2 mL/L), uracil (56 mg/L), uridine (122.1 mg/L), and riboflavin (2.5 mg/L). The trace elements stock solution (500X) contained FeSO_4_·7H_2_O (1 g/L), ZnSO_4_·7H_2_O (8.8 g/L), CuSO_4_·5H_2_O (0.4 g/L), MnSO_4_·H_2_O (0.15 g/L), Na2B4O7·10H_2_O (0.1 g/L), and (NH_4_)_6_Mo_7_O_24_·4H_2_O (0.05 g/L). Regular MM contained 1% final D+ glucose (10 g/L), MgSO4 (2 mL/L of 26% w/v), trace elements solution (2 mL/L), and stock salt solution (10 mL/L). Sorbose MM, to induce hyphal tip lysis^41^, contained 2% final L-sorbose (20 g/L), 0.1% final D+ glucose (1 g/L), trace elements solution (2 mL/L), and stock salt solution (10 mL/L). The stock salt solution (100x; pH 6-6.5) contained NaNO_3_ (60 g/L), KCl (5.2 g/L), and KH_2_PO_4_ (15.2 g/L). For strains with auxotrophic alleles, MM was supplemented as follows: *pyrG89* allele: 0.5 mM uracil and 0.5 mM uridine; *pabaA1* allele: 1.46 μM p-aminobenzoic acid; *riboB2* allele: 6.6 μM riboflavin; *pyroA4* allele: 3.0 μM pyridoxine.

To grow adult hyphae for imaging, spores were inoculated into MM agar with supplements and incubated for 16-20hr at 37°C. Individual colonies were cut from the agar and inverted onto 35 mm #1.5 glass-bottom imaging dishes (Cellvis D35C4201.5N). To grow germlings for imaging, spores were collected into 0.01% Tween-80 (Sigma-Aldrich P1754) and their density was measured using a hemocytometer. The spores were then inoculated into liquid MM with supplements at a density of 5×10^5^ spores/mL in a four-chamber 35 mm dish and incubated for 16-20 hr at 30°C in an unsealed humid plastic container.

### Phylogenetic analysis of peroxisome hitchhiking proteins

Pezizomycotina fungi included in this analysis were previously published to have Woronin bodies and/or have been referenced as representatives of different clades^42,43^. Species were only included if they had their whole genome sequenced and proteome annotated and/or drafted at the time of this analysis (30 August 2022). The phylogenetic tree for fungi was drawn manually based on the taxonomic rankings of each species on NCBI.

Homologous genes for PxdA (AN1156) and DipA (AN10946) were identified using BLASTp available at NCBI (https://blast.ncbi.nlm.nih.gov/) (parameters: E = 1.0×10^−5^, word size of 6, BLOSUM62 matrix, gap costs existence 11 and extension 1, filtering low complexity regions, minimum percent coverage 20%). Reciprocal BLASTs were used to confirm all candidate homologs. The reference proteins database (refseq_proteins) was used to reduce redundancy of search results. For species where the refseq_proteins database was unavailable, the non-redundant protein sequences database (nr) was used instead. The E value, query coverage, and percent identity of the top hits for a reference strain of each species was recorded (Table S1).

To identify PxdA homologs, three query sequences were used: the PxdA coiled-coil region, previously defined as CC1-3 (aa 1466-2115)^7^; the PxdA uncharacterized N-terminus (aa 1-1465); and the full-length PxdA protein (aa 1-2236). The conserved stretch of amino acids in the PxdA CC3 domain are aa 2033-2058. The coiled-coil region of all PxdA homologs was individually predicted by querying each sequence in the Waggawagga coiled-coil prediction software (https://waggawagga.motorprotein.de/) with the following settings: Marcoil, Scorer 2.0 oligomerization, and window length of 21. PxdA possesses a predicted coiled-coil region that is necessary and sufficient for peroxisome hitchhiking^7^. The coiled-coil prediction software Waggawagga estimated between three to six distinct coiled-coils in each PxdA homolog, with the total coiled-coil region varying from 167 to 307 aa long (Table S1). All PxdA homologs possessed an N-terminal extension of variable length with no discernable conserved domains, based on NCBI and PFAM databases.

For identifying DipA homologs, query sequences included both the full-length DipA protein (aa 1-704) and the DipA metallophosphatase domain alone (aa 45-96). DipA homologs were defined to have both a metallophosphatase domain followed by a conserved domain of unknown function (DUF2433). The amino acid boundaries of both the metallophosphatase domains and the DUF2433 were identified for all DipA homologs using NCBI domain prediction and MUSCLE multiple protein sequence alignments to the *A. nidulans* DipA reference sequence in SnapGene.

### Spinning disk confocal microscopy

Live cell imaging was performed at room temperature using a Nikon Ti2 microscope mounted with a Yokogawa W1 spinning-disk confocal scan head and two Prime95B cameras (Photometrics). All images were acquired in 16-bit format. The NIS Elements Advanced Research software (Nikon) was used to run the microscope using the 488 nm and 561 nm lasers of a six-line LUN-F-XL laser engine (405 nm, 445 nm, 488 nm, 515 nm, 561 nm, and 640 nm). The stage xy position was controlled by a ProScan linear motor stage controller (Prior) and its z position was controlled by a piezo Nano-Z stage positioner (Mad City Labs). An Apo TIRF 100x 1.49 NA objective (Nikon) was used for all experiments except for the imaging of apical septa for hyphae grown in glucose and sorbose agar, which required an S Fluor 40x 1.30 NA objective (Nikon) to capture a larger field of view.

Simultaneous dual-color imaging was used to assess colocalization of Woronin body and peroxisome proteins in germlings. Z-stacks were collected to capture the complete volume of germling cells in the field of view (0.2 μm step size, 8-10 μm total depth). The 488 nm and 561 nm lines were used to excite mGFP5- and mCherry-tagged fusion proteins, respectively. The emission was split with a Cairn TwinCam with a 580LP filter. The GFP/488 emission was reflected and passed through a 514/30 bandpass filter onto camera two. The mCherry/561 fluorescence was passed through a 617/73 and an additional 600/50 bandpass filter to camera one. The cameras were aligned manually in NIS Elements using 0.1 μm Tetra-Speck beads (Thermofisher T14792). Image acquisition settings, such as exposure time and laser power, were determined at the beginning of each imaging session to eliminate fluorescent bleed through between each channel.

Triggered acquisition of 488/561 was used when imaging Woronin body association at apical septa with the 40x objective and when imaging SspA-mGFP5 comigration with PxdA-mKate with the 100x objective. The firing of the 488 nm and 561 nm lasers was synchronized by the Prime95B camera trigger signal, which was integrated into a Nikon BB connected to a 6723 DAQ board housed in an external Pxi chassis (National Instruments). A quad bandpass filter (Chroma ZET405/488/561/640mv2) was placed in the emission path of the W1 scan head.

### Quantitative image analysis

All image analysis was performed using ImageJ/FIJI (National Institutes of Health, Bethesda, MD). All microscopy images were blinded prior to analysis using a custom script in R^44^. Statistical tests were performed using GraphPad Prism (Windows version 9.4.1). *Code availability:* All macros (ImageJ, Windows version 1.53q) and data processing scripts (R, Windows version 4.1.0; Rstudio, Windows version 2022.12.0 Build 353) are available on GitHub (https://github.com/LiviaSongster).

### Colocalization and co-occurrence

To prepare Z-stacks for colocalization analysis, whole germlings were cropped, max-Z projected, and saved as separate files. For both the 488 and 561 channel, a Gaussian Blur (sigma = 1) and background subtraction (rolling = 50) were applied prior to thresholding (Yen dark) to generate a binary puncta mask. The threshold for the green channel was set at 15 to 65535, and for the red channel at 20 to 65535. All gray values outside of that channel’s mask were cleared to eliminate background. Colocalization was then quantified using Pearson’s coefficient of correlation and Mander’s coefficient of co-occurrence algorithms via the JACoP plugin in ImageJ^45,46^.

The fraction of SspA puncta with PTS1 and the fraction of SspA puncta with PTS1 was manually scored for four separate experiments containing at least ten germlings each. Germlings were imaged for 60 sec (300 ms interval) with a single z-plane using simultaneous dual-color settings. Puncta in the cytoplasm or at septa were manually segmented as an ROI and scored as colocalized if there was a punctum from both channels that overlapped by >50% and if the mean gray value of the puncta of the opposite channel was at least 10% brighter than the background. Motile puncta were assessed by first generating kymographs, and for directed runs where a punctum traveled >2 μm, the co-occurrence of puncta in both channels was manually scored.

To measure the fluorescence intensity ratio of different proteins, puncta from sum Z projections of germlings were manually circled as regions of interest (ROIs). The gray value of all puncta was measured in both the 488 and 561 channel and the intensity ratio for each ROI was calculated as the mean gray value of the 488 channels divided by the mean gray value of the 561 channels.

### Organelle flux

Total puncta movement was estimated from single-plane, triggered-acquisition movies (30 sec total length, 300 ms interval, 100 ms exposure per laser). The number of puncta crossing a perpendicular line approximately 10 μm from the hyphal tip was manually counted.

### Intensity profile during movement

The intensity along a line (1 pixel width) was measured during a directed run for both the 488 and 561 channel. The line scans from multiple kymographs were compiled in R. For each line scan, the maximum intensity in the 561 channel was set as x position 0 μm and then applied to the 488 channel.

### Speed and distance travelled of migrating puncta

Adult hyphae were imaged for 60 sec (300 ms interval) with a single z-plane using simultaneous dual-color settings. The line tool was used to trace puncta trajectories over time and movies were resliced along these lines to generate kymographs. The instantaneous speed and run lengths of individual moving puncta were calculated from the inverse of the slopes of manual kymograph traces. A punctum was scored as moving if it exhibited a directed run (no pauses) > 3 μm during its movement.

### Hyphal tip line scans

Whole adult hyphae were imaged live as a Z-stack (0.2 μm step size, 8-10 μm total depth). Max-Z intensity projections of fluorescence micrographs were obtained and the brightfield channel was used to trace from the hyphal tip inwards using the segmented line tool (pixel width 20). The traces were superimposed on the fluorescence micrographs to project the average fluorescence intensity along the line. The mean background intensity was measured for each cell and subtracted from that cell’s respective line scan prior to binning.

### Woronin body localization at tip apical septa

Cells expressing mCherry-SsoA and SspA-mGFP5 were grown overnight with supplements in either regular MM (1% glucose) or sorbose MM (2% sorbose/0.1% glucose). Adult hyphae were imaged using the 40x objective to collect complete volume Z-stacks (0.2 μm step size, 8-10 μm total depth). Images were max-Z projected and the apical septa were manually identified as ROIs and cropped. Crops of single septa were blinded and Woronin body localization was manually scored as the fraction of apical septa with an associated Woronin body per experiment. The mean fluorescence intensity of SspA-mGFP5 from these max-Z projections was measured within a 0.5 μm-wide by 1 μm-long rectangle centered at an apical septum. The mean background intensity adjacent to each septum was measured and subtracted from the mean septal intensity. Background-corrected measurements were then normalized to the average septal intensity of wild type.

### Cytoplasmic leakage

Cytoplasmic leakage was measured for burst hyphal tips in sorbose agar. First, complete Z stacks (0.2 μm step size, 4 μm total depth) were imaged, sum-Z projected, and blinded. The integrated density (mean intensity of puddle times puddle area) for each cytoplasmic puddle was measured in ImageJ and corrected by subtracting the product of the mean background intensity and puddle area. This background-corrected integrated density was then normalized to the mean observation for wild type to calculate the normalized amount of leaked cytoplasm.

As a complimentary approach, mycelia were also subjected to hypotonic shock to induce global tip lysis, using an adapted protocol for *A. nidulans*^47^. Spores from strains expressing a cytoplasmic GFP were inoculated into liquid YG media and grown overnight at 37°C. Mycelia was collected in Miracloth (EMD Millipore 475855) and dried with paper towels. Approximately 300 ± 10 mg dried mycelia were submerged in 1 mL sterile ddH_2_O and vortexed to induce hypotonic damage at hyphal tips. Samples were spun at 15,000 x g for 1 min in a tabletop centrifuge to pellet cell debris and the supernatant was diluted 1:10 in sterile ddH_2_O for a 96-well microplate Bradford microassay (Bio-Rad 5000201) to estimate the amount of leaked protein. The absorbance at 595 nm was measured using a Cytation 5 plate reader (BioTek) and Gen5 data analysis software (Windows version 3.05).

**Figure S1.**
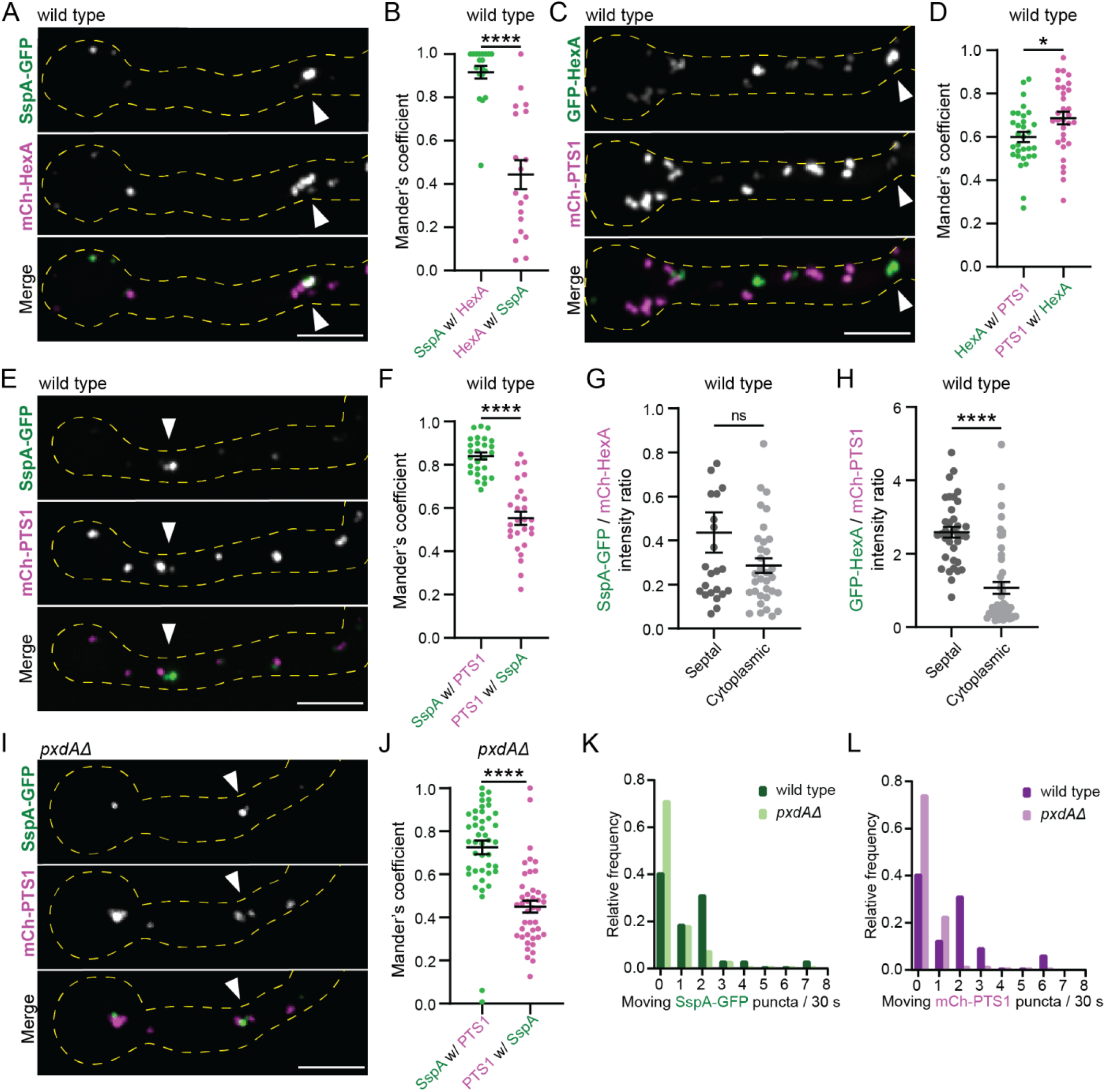
Woronin body and peroxisome colocalization and co-occurrence in whole germlings. **(A, C, E, I)** Example max-Z projections of colocalization between GFP and mCherry-tagged markers for Woronin bodies and peroxisomes in wild type (A, C, E) or *pxdAΔ* (I) germlings. White arrows indicate the septum closest to the spore head. Scale bars = 5 μm. **(B, D, F)** Average Mander’s coefficient for each pair of markers in wild type germlings. B) SspA-mGFP5 and mCherry-HexA puncta in wild type (n = 19 germlings; mean ± SD: SspA w/ HexA = 0.92 ± 0.13; HexA w/ SspA = 0.44 ± 0.29). D) mGFP5-HexA and mCherry-PTS1 puncta in wild type (n = 31 germlings; mean ± SD: HexA w/ PTS1 = 0.60 ± 0.13; PTS1 w/ HexA = 0.69 ± 0.16) F) SspA-mGFP5 and mCherry-PTS1 puncta in wild type (n = 27 germlings; mean ± SD: SspA w/ PTS1 = 0.84 ± 0.085; PTS1 w/ SspA = 0.55 ± 0.16). **(G)** Average intensity ratio of SspA-mGFP5 and mCherry-HexA in puncta at septa and in the cytoplasm (n puncta, mean ± SD: septal = 24, 0.44 ± 0.45; cytoplasmic = 33, 0.29 ± 0.19). **(H)** Average intensity ratio of mGFP5-HexA and mCherry-PTS1 in puncta at septa and in the cytoplasm (n, mean ± SD: septal = 36, 2.59 ± 0.89; cytoplasmic = 47, 1.1 ± 1.1). **(J)** SspA-mGFP5 and mCherry-PTS1 puncta in *pxdAΔ* germlings (n = 43 germlings; mean ± SD: SspA w/ PTS1 = 0.73 ± 0.21; PTS1 w/ SspA = 0.45 ± 0.18). **(K-L)** Histograms of the relative frequency of all movements (retrograde and anterograde) for SspA-mGFP5 (K) and mCherry-PTS1 (L) in adult hyphae (n hyphal tips: wild type = 32; *pxdAΔ* = 64). **For B, D, F-H, and J:** unpaired t-test. Error bars represent SEM. P > 0.05 (labeled ns for “not significant,” or not shown), P ≤ 0.05 (*), P ≤ 0.01 (**), P ≤ 0.001 (***), P ≤ 0.0001 (****).

**Figure S2.**
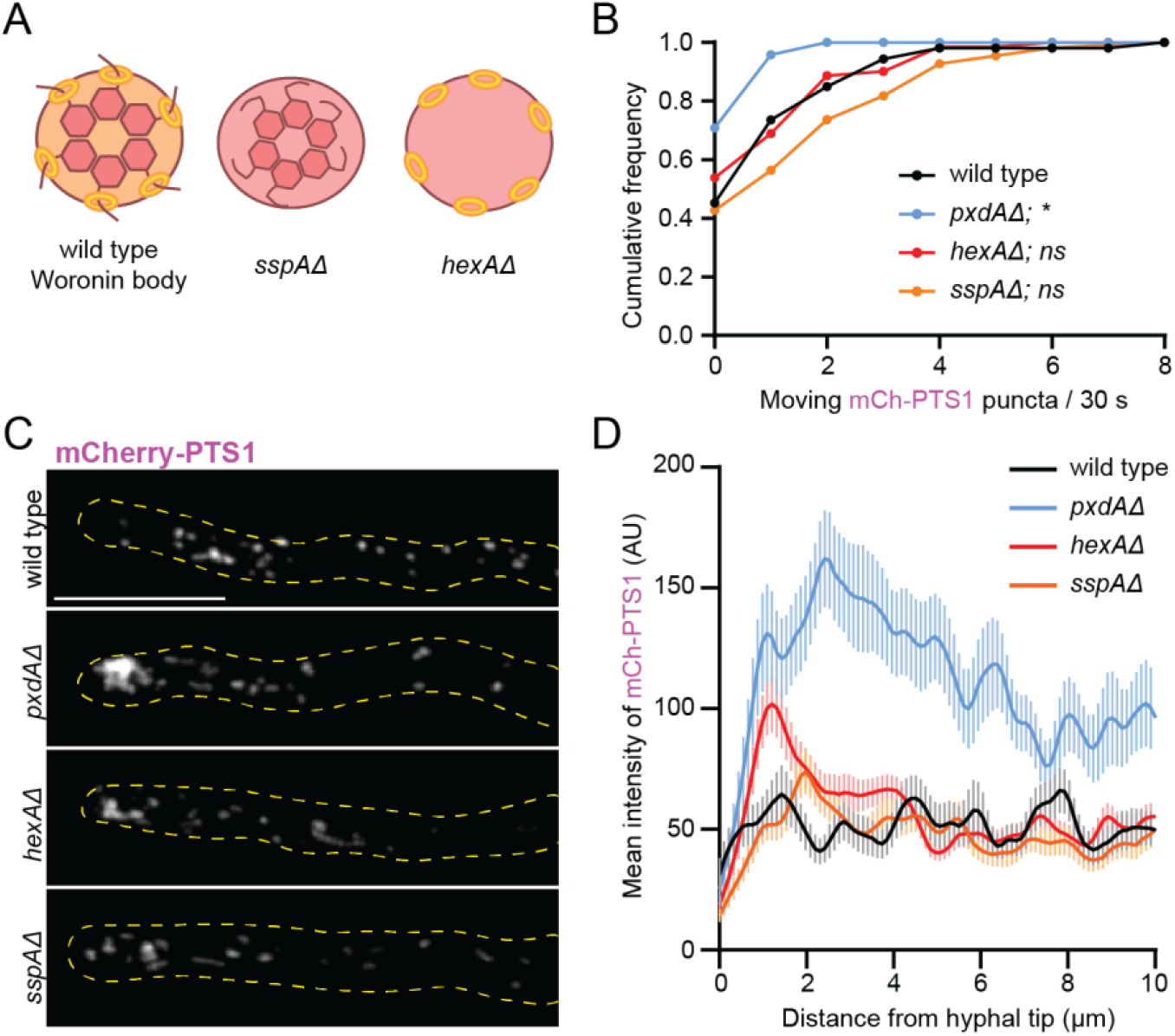
Peroxisome distribution and movement in cells lacking complete Woronin bodies. **(A)** Diagram of Woronin bodies lacking SspA or HexA. **(B)** Cumulative frequency plot of the number of moving mCherry-PTS1 puncta per 30 s in adult hyphal tips (n, mean ± SD puncta/30 s: wild type = 53 hyphae, 1.09 ± 1.5; *pxdAΔ* = 48 hyphae, 0.33 ± 0.56; *hexAΔ* = 132 hyphae, 1.02 ± 1.4; *sspAΔ* = 110 hyphae, 1.60 ± 1.86). The results of a Kruskal-Wallis with post-hoc Dunn’s multiple comparisons test versus wild type are shown; P > 0.05 labeled ns for “not significant”; P ≤ 0.05 (*). **(C)** Examples of mCherry-PTS1 in adult hyphal tips (max-Z projections). Scale bar = 10 μm. **(D)** Mean intensity of mCherry-PTS1 from the hyphal tip inwards (n: wild type = 33 hyphae; *pxdAΔ* = 30 hyphae; *hexAΔ* = 53 hyphae; *sspAΔ* = 54 hyphae). Error bars represent SEM.

**Figure S3.**
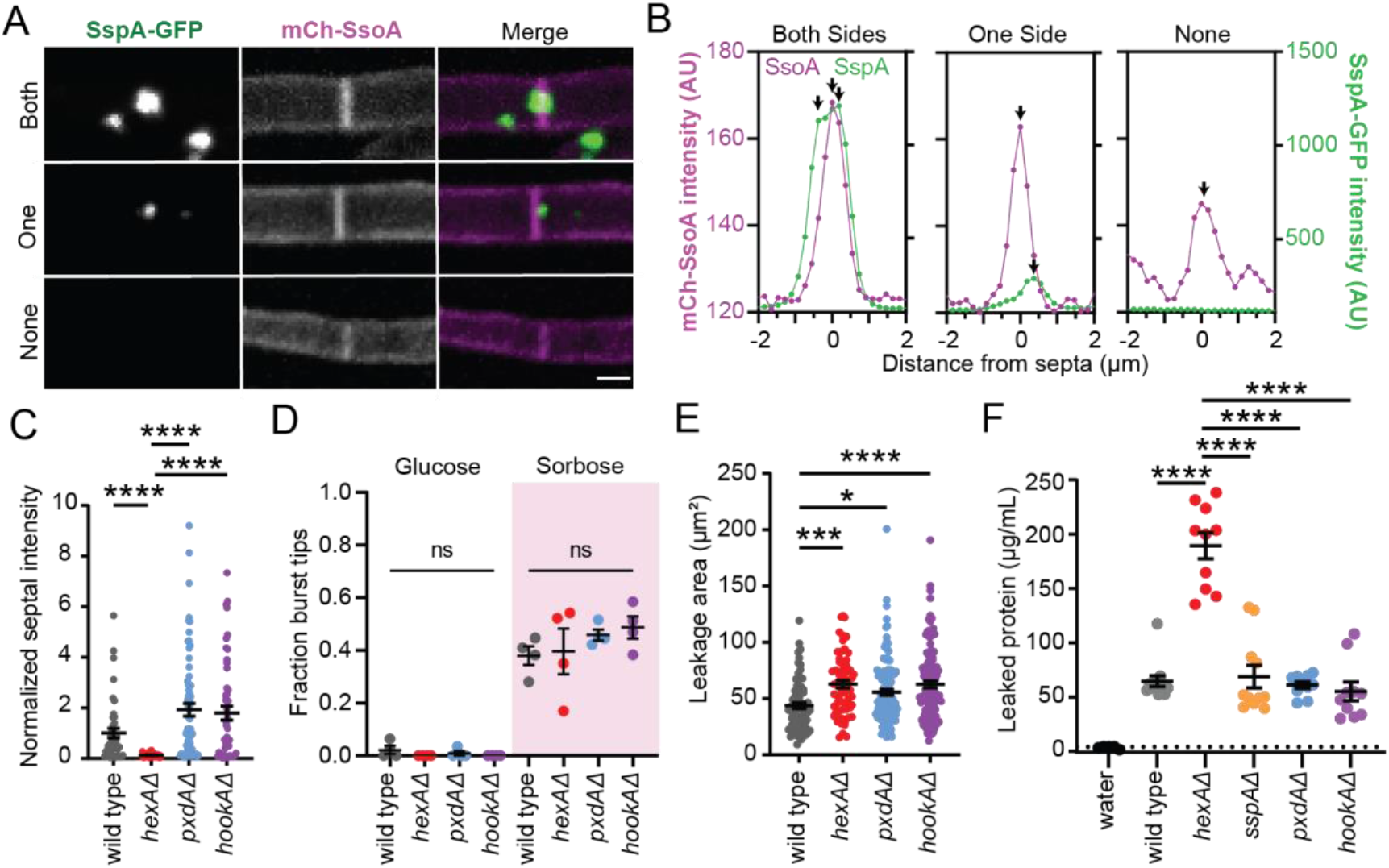
Woronin bodies at tip apical septa and cytoplasmic leakage assays. **(A)** Examples of Woronin body localization at septa. SspA-mGFP5 puncta were scored on max-Z projections as either present on both sides, one side, or none. Scale bar = 2 μm. **(B)** Line scans showing fluorescence intensity of Woronin bodies (SspA) or cell membrane at septa (SsoA) for each score category from A. Arrows indicate peak intensities for each fluorescent protein. **(C)** Normalized mean intensity of SspA-mGFP5 on either side (±0.5 μm) of apical septa for cells grown in glucose MM agar (n septa, mean ± SD AU: wild type = 43, 1.00 ± 1.24; *hexAΔ* = 45, 0.12 ± 0.6; *pxdAΔ* = 68, 1.93 ± 2.08; *hookAΔ* = 50, 1.80 ± 1.98). **(D)** Fraction of burst or damaged hyphal tips when grown in glucose or sorbose agar. Each circle represents the fraction burst hyphae from all tips imaged in a single experiment with 8-50 hyphae each (Glucose n experiments, mean ± SD: wild type = 4, 0.02 ± 0.03; *hexAΔ* = 4, 0 ± 0; *pxdAΔ* = 4, 0.01 ± 0.02; *hookAΔ* = 4, 0 ± 0. Sorbose n, mean ± SD: wild type = 4, 0.38 ± 0.07; *hexAΔ* = 4, 0.39 ± 0.17; *pxdAΔ* = 4, 0.46 ± 0.04; *hookAΔ* = 4, 0.49 ± 0.08. **(E)** Leakage area of cytoplasmic puddles for hyphae grown in sorbose (n burst hyphae, mean ± SD: wild type = 73, 43.7 ± 22.7 μm^2^; *hexAΔ* = 63, 62.7 ± 26.8 μm^2^; *pxdAΔ* = 125, 55.6 ± 28.9 μm^2^; *hookAΔ* = 119, 62.5 ± 32.5 μm^2^). **(F)** Quantification of cytoplasmic leakage after hypotonic shock. The amount of leaked protein in the supernatant was quantified via Bradford assay (n samples, mean ± SD: water = 5, -0.45 ± 1.3 μg/mL; wild type = 12, 62 ± 18 μg/mL; *hexAΔ* = 10, 189 ± 39 μg/mL; *sspAΔ* = 11, 66 ± 35 μg/mL; *pxdAΔ* = 10, 58 ± 28 μg/mL; *hookAΔ* = 10, 52 ± 28 μg/mL). **For C:** Kruskal-Wallis test with Dunn’s multiple comparisons for nonparametric data. **For D-F:** Ordinary one-way ANOVA with post-hoc Tukey-Kramer test. **For C-F:** Error bars represent SEM. P > 0.05 (labeled ns for “not significant,” or not shown), P ≤ 0.05 (*), P ≤ 0.01 (**), P ≤ 0.001 (***), P ≤ 0.0001 (****).

**Movie S1 (separate file)**. Woronin body (SspA-mGFP5) and peroxisome (mCherry-PTS1) comigration in a wild type hyphal tip. A single Z plane image was taken every 300 ms via simultaneous dual-color spinning disk confocal microscopy (Nikon). The white arrow indicates a comigrating, bidirectional run. Scale bar = 10 μm.

**Movie S2 (separate file)**. Woronin bodies (SspA-mGFP5) and peroxisomes (mCherry-PTS1) in a *pxdAΔ* hyphal tip. A single Z plane image was taken every 300 ms via simultaneous dual-color spinning disk confocal microscopy (Nikon). Scale bar = 10 μm.

**Table S1 (separate file)**. PxdA (sheet 1) and DipA (sheet 2) BLASTp results for all 33 fungal species tested. The amino acid sequences used for each BLASTP search are shown in row 1 on both sheets.

**Table S2.**
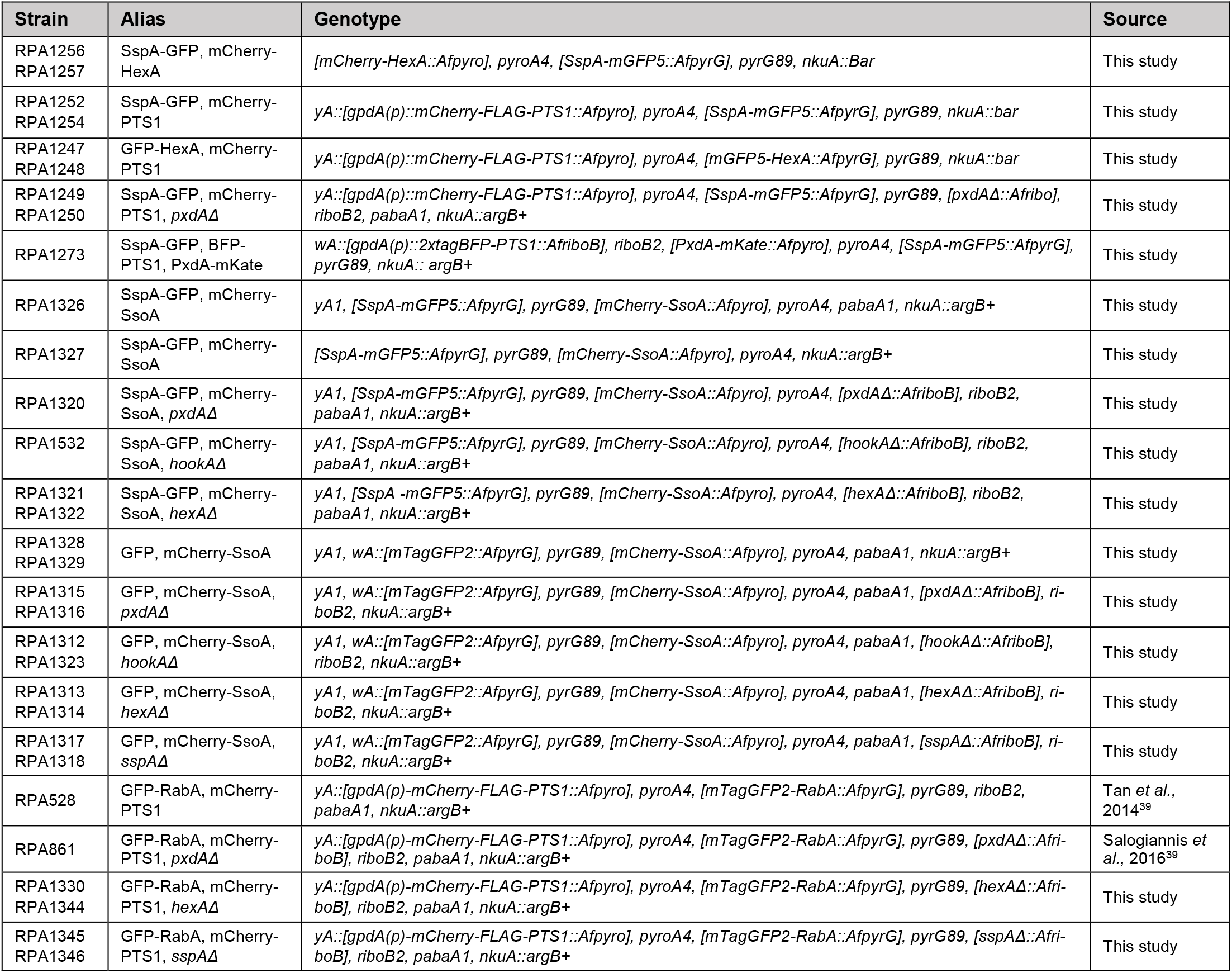
Strains used in this study.

**Table S3.**
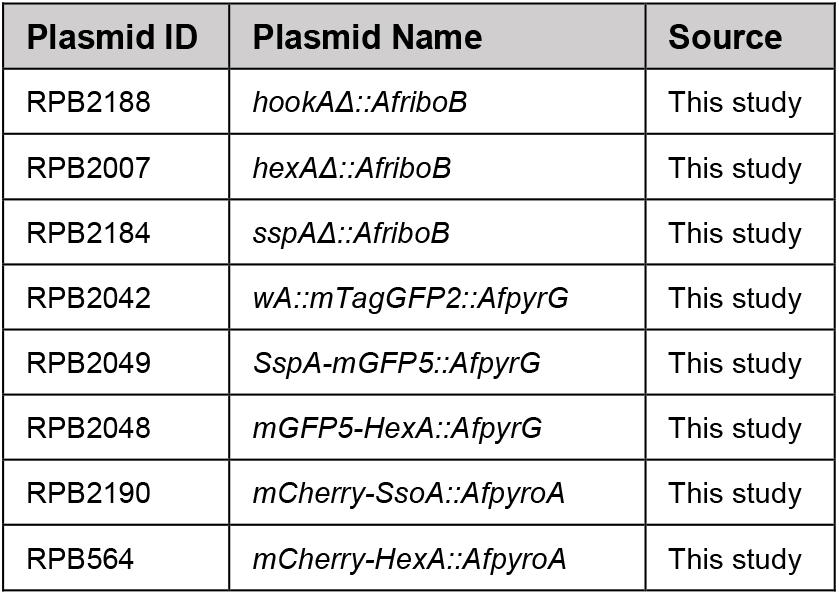
Plasmids generated for this study.

**Table S4.**
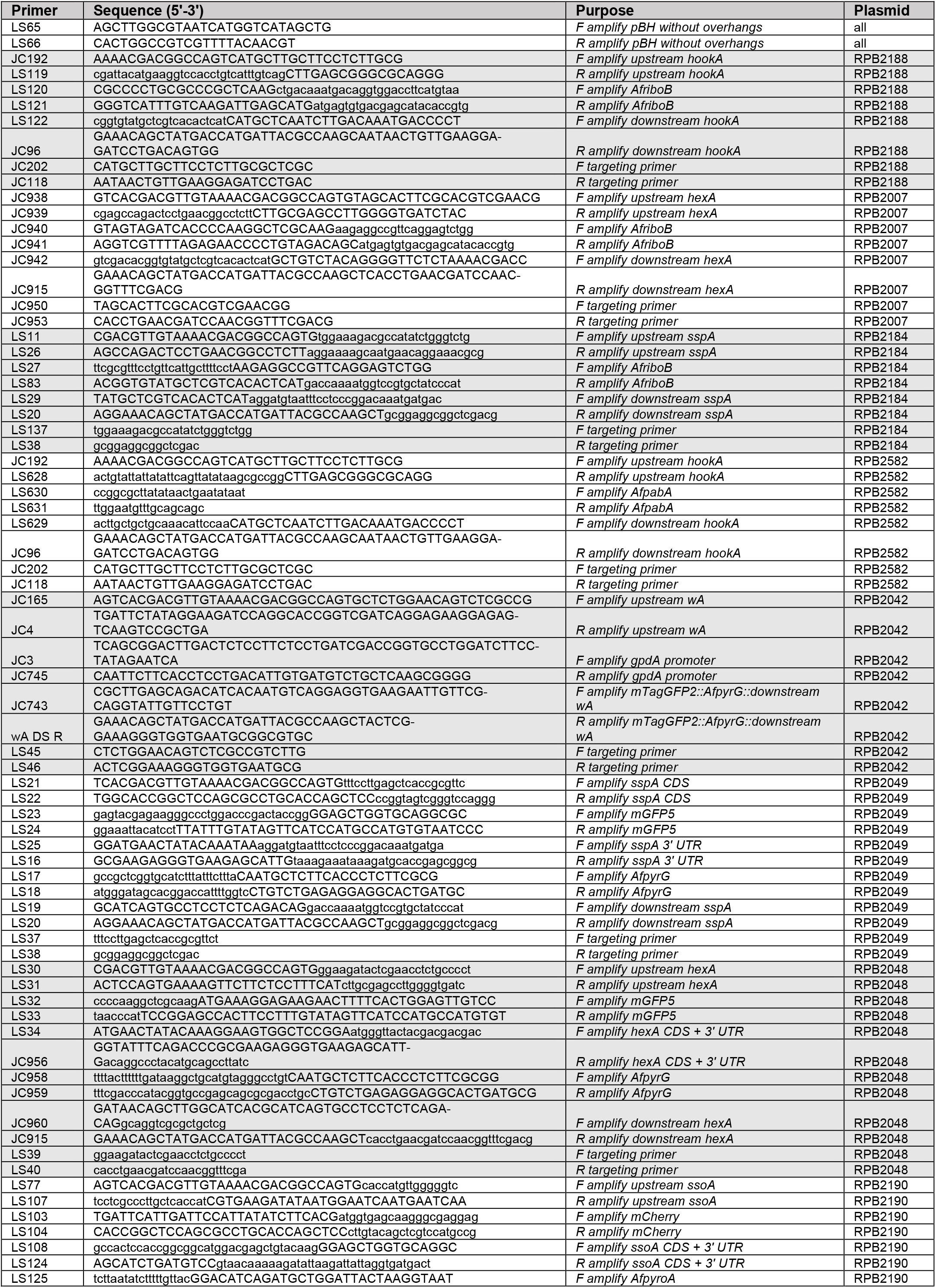
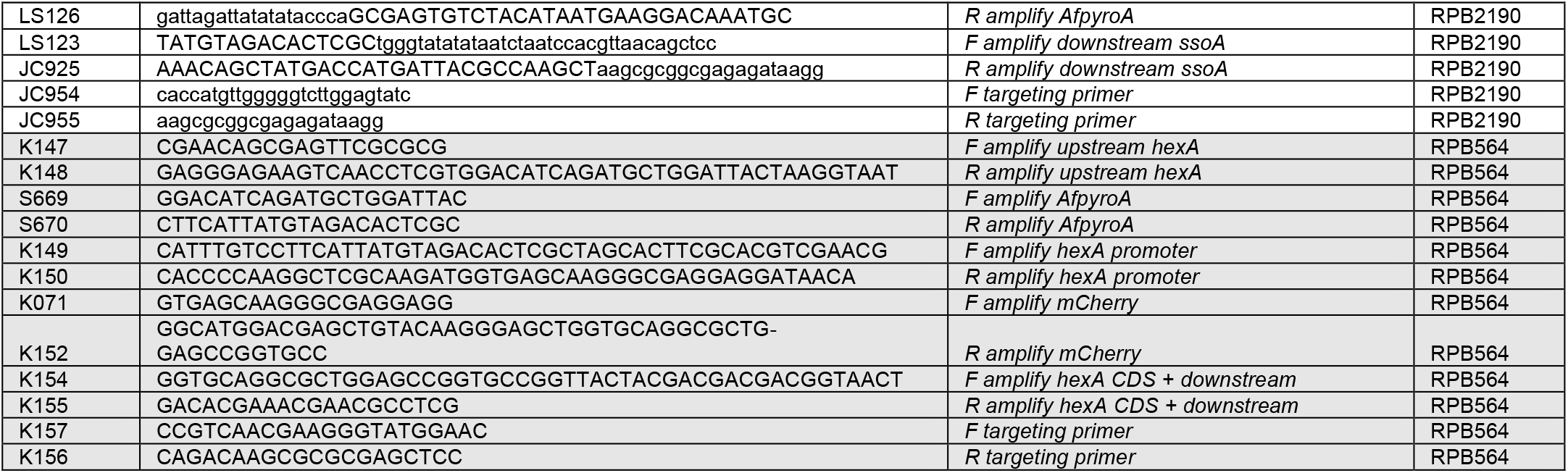
Primers used to make plasmids for this study.

